# Cognitive ability and physical health: a Mendelian randomization study

**DOI:** 10.1101/084798

**Authors:** Saskia P Hagenaars, Catharine R Gale, Ian J Deary, Sarah E Harris

## Abstract

Causes of the association between cognitive ability and health remain unknown, but may reflect a shared genetic aetiology. This study examines the causal genetic associations between cognitive ability and physical health. We carried out two-sample Mendelian randomization analyses using the inverse-variance weighted method to test for causality between later life cognitive ability, educational attainment (as a proxy for cognitive ability in youth), BMI, height, systolic blood pressure, coronary artery disease, and type 2 diabetes using data from six independent GWAS consortia and the UK Biobank sample (N = 112 151). BMI, systolic blood pressure, coronary artery disease and type 2 diabetes showed negative associations with cognitive ability; height was positively associated with cognitive ability. The analyses provided no evidence for casual associations from health to cognitive ability. In the other direction, higher educational attainment predicted lower BMI, systolic blood pressure, coronary artery disease, type 2 diabetes, and taller stature. The analyses indicated no causal association from educational attainment to physical health. The lack of evidence for causal associations between cognitive ability, educational attainment, and physical health could be explained by weak instrumental variables, poorly measured outcomes, or the small number of disease cases.

## Introduction

Lower cognitive ability, lower educational attainment and greater cognitive decline are all associated with poorer health outcomes ^1–3^. Some of these associations possibly arise because of the effect of lower cognitive ability in childhood on later life health, others because illnesses may lower cognitive ability in later life. The causes of these associations are unclear, but some may reflect, in part, a shared genetic aetiology. Recent papers have reported genetic associations between cognitive ability and educational attainment, and a number of physical and mental health traits and diseases ^4–6^. These ^4,6^, and other papers^7–9^, have shown successful use of educational attainment as a proxy for cognitive ability, showing phenotypic correlations between educational attainment and general cognitive ability around 0.50 ^9^ and a genetic correlation of 0.72 ^4^.

Some of the reciprocal phenotypic associations between cognitive and physical health variables, and their genetic correlations, are as follows. Short stature has been consistently linked with lower cognitive ability ^10,11^. Molecular genetic studies have indicated positive genetic correlations between height and cognitive ability ^4,12^, as well as between height and educational attainment ^4,5^. Higher polygenic scores for height have been associated with better cognitive ability in adulthood ^4^. A causal association was reported between taller stature and educational attainment (not including individuals with a degree) in UK Biobank using a Mendelian randomization analysis^13^.

Multiple studies have shown associations between cognitive ability and cardiovascular risk factors. For example, lower childhood cognitive ability is associated with subsequent high blood pressure ^14^ and obesity ^15^. However, higher BMI in mid-life ^16^ and both hypertension and hypotension^17^ are associated with lower cognitive ability and greater cognitive decline in later life. A negative genetic correlation has been identified between BMI, but not blood pressure, and educational attainment and cognitive ability in mid to late life ^4,5^, and a polygenic score for higher BMI is associated with lower cognitive ability in mid to late life and lower educational attainment ^4^; however, a polygenic score for higher systolic blood pressure is associated with lower educational attainment, but higher cognitive ability in mid to late life ^4^.

Similarly, associations have been identified between cognitive ability and cardio metabolic diseases. Childhood cognitive ability has been associated with developing diabetes ^18^ and coronary artery disease ^19^ later in life. Diabetes ^20^ and coronary artery disease ^21,22^ in midlife have been associated with greater cognitive decline later in life. A polygenic risk score for type 2 diabetes is associated with lower educational attainment, but not with cognitive ability in mid to late life ^4^, although one has been associated with reduced cognitive decline ^23^. To date, no significant genetic correlation between diabetes and cognitive ability has been identified ^4,5^. A polygenic risk score for coronary artery disease is associated with lower educational attainment and lower mid to late life cognitive ability ^4^, and a negative genetic correlation was identified between coronary artery disease and educational attainment ^4,5^, but not cognitive ability in mid to late life ^4^.

The question arises as to whether the genetic cognitive-health associations caused by: 1) genes influencing health traits/diseases, and then those health traits/diseases subsequently influencing cognitive ability; 2) genes influencing cognitive ability, and then cognitive ability subsequently influencing health traits/diseases; 3) genes influencing general bodily system integrity^24^ that influences both cognitive ability and health traits/diseases?

To try to make some progress in understanding causality of the correlation between cognitive ability and a number of physical and mental health traits, in the present report we used a bidirectional, two-sample Mendelian randomization (MR) approach ^25^. MR uses genetic variants as proxies for environmental exposures and is subject to the following assumptions: 1) the genetic variants are associated with the exposure; 2) the genetic variants are only associated with the outcome of interest via their effect on the exposure [i.e., there is no biological pleiotropy (the phenomenon whereby one SNP independently influences multiple traits), also called the exclusion restriction]; and 3) the genetic variants are independent of confounders. Figure 1 shows the Mendelian randomization study model; the instrumental variable, here based on genome-wide significant SNPs from independent studies for the exposure, is used to estimate if the exposure (e.g. BMI) causally influences the outcome (e.g. cognitive ability). Individual single nucleotide polymorphisms (SNPs) are often found to be weak instruments for investigating causality because they often have small effect sizes. Using multiple SNPs can increase the strength of the instrument. However, this increases the chance of violating the MR assumptions, specifically violation of the assumption that the genetic variants affect the outcome only via the exposure. We used multiple genetic variants for a number of health-related traits and diseases, previously identified in genome-wide association studies, as instrumental variables to see if they predicted cognitive ability (verbal-numerical reasoning) in mid to later life in the UK Biobank. We then used genome-wide significant educational attainment SNPs as an instrumental variable to test whether genetic differences associated with educational attainment (a proxy measure of cognitive ability in early life ^6,8^) predict later life health outcomes in the UK Biobank.

**Figure 1.**
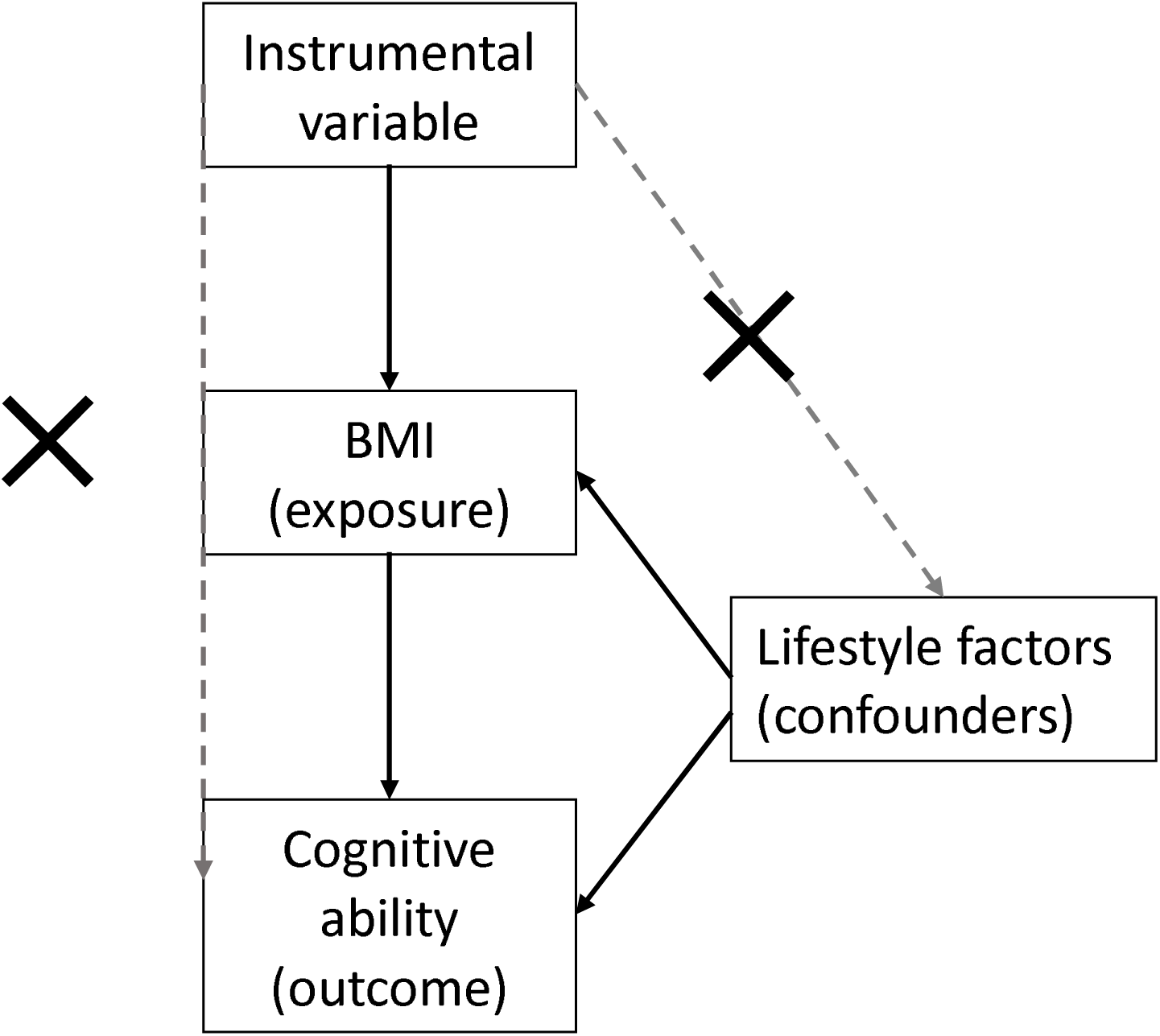
Model for Mendelian randomization study. The instrumental variable, based on genome-wide significant SNPs from independent studies for the exposure, is used to estimate if the exposure (e.g. BMI) causally influences the outcome (e.g. cognitive ability). The instrumental variable should be unrelated to potential confounders of the exposure-outcome association and should only affect the outcome via the exposure.

## Methods

### Sample

This study uses baseline data from the UK Biobank Study, a large resource for identifying determinants of human diseases in middle aged and older individuals ^26^. Around 500 000 community-dwelling participants aged between 37 and 73 years were recruited and underwent assessments between 2006 and 2010 in the United Kingdom. This included cognitive and physical assessments, providing blood, urine and saliva samples for future analysis, and giving detailed information about their backgrounds and lifestyles, and agreeing to have their health followed longitudinally. For the present study, genome-wide genotyping data were available on 112 □ 151 individuals (58 □ 914 females) aged 40-70 years (mean age=56.9 years, SD=7.9) after the quality control process which is described in more detail elsewhere ^4^. The UK Biobank study was approved by the National Health Service (NHS) Research Ethics Service (approval letter dated 17th June 2011, reference: 11/NW/0382). The analyses in the present report were completed under UK Biobank application 10279. All experiments were performed in accordance with guidelines and regulations from these committees. Written informed consent was obtained from each subject.

## Measures

### Body mass index

Body mass index (BMI) was calculated as weight(kg)/height(m)^2^, and measured using an impedance measure, i.e. a Tanita BC418MA body composition analyser, to estimate body composition. We used the average of the two methods when both measures were available (r = 0.99); if only one measure was available, that measure was used (N = 1629). 291 individuals did not have information on BMI. One outlier was excluded based on visual inspection of the BMI distribution (BMI > 50). 111 712 individuals had valid BMI and genetic data.

### Height

Standing and sitting height (cm) were measured using a Seca 202 device. We used standing height and excluded one individual based on the visual inspection of the height distribution with a standing height < 125 cm and a sitting/standing height ratio < 0.75. 111 959 had valid height and genetic data.

### Systolic blood pressure

Systolic blood pressure was measured twice, a few moments apart, using the Omron Digital blood pressure monitor. A manual sphygmomanometer was used if the digital blood pressure monitor could not be employed (N = 6652). Systolic blood pressure was calculated as the average of measures at the two time points (for either automated or manual readings). Individuals with a history of coronary artery disease were excluded from the analysis (N = 2513). Following the recommendation by Tobin, et al. ^27^, 15 mmHg was added to the average systolic blood pressure of individuals taking antihypertensive medication (N = 10 988). Individuals with a systolic blood pressure (after correcting for medication) more than 4 SD from the mean were excluded from future analyses (N = 75). After all exclusions, 106 759 individuals remained with valid blood pressure and genetic data.

### Coronary artery disease

UK Biobank participants completed a touch screen questionnaire on past and current health, which included the question “Has a doctor ever told you that you have had any of the following conditions? heart attack/angina/stroke/high blood pressure/none of the above/prefer not to answer”. This was followed by a verbal interview with a trained nurse who was made aware if the participant had a history of certain illnesses and confirmed these diagnoses with the participant. For the present study, coronary artery disease was defined as a diagnosis of myocardial infarct or angina, reported during both the touchscreen and the verbal interview in individuals with genetic data (N = 5288). The control group (N = 104 784) consisted of participants who reported none of the following diseases (based on the non-cancer illness code provided by UK Biobank): myocardial infarction, angina, heart failure, cerebrovascular disease, stroke, transient ischaemic attack, subdural haemorrhage, cerebral aneurysm, peripheral vascular disease, leg claudication/intermittent claudication, arterial embolism.

### Type 2 diabetes

Type 2 diabetes case-control status was created using the same method as described by Wood, et al. ^28^, for all individuals with genetic data based on the interim release of UK Biobank. Cases included participants who reported type 2 diabetes or generic diabetes during the nurse interview, started insulin treatment at least one year after diagnosis, were older than 35 years at the time of diagnosis, and did not receive a diagnosis one year prior to baseline testing (N = 3764). The control group consisted of participants who did not fulfil these criteria, and did not report a diagnosis of type 1 diabetes, diabetes insipidus and gestational diabetes (N = 108 015).

### Years of education

As part of the sociodemographic questionnaire in the study, participants were asked, “Which of the following qualifications do you have? (You can select more than one)”. Possible answers were: “College or University Degree/A levels or AS levels or equivalent/O levels or GCSE or equivalent/CSEs or equivalent/NVQ or HND or HNC or equivalent/Other professional qualifications e.g. nursing, teaching/None of the above/Prefer not to answer”. For the present study, a new continuous variable was created measuring ‘years of education completed’. This was based on the ISCED coding, using the 1997 International Standard Classification of Education (ISCED) of the United Nations Educational, Scientific and Cultural Organization ^29^. See the Table 1 for further details. Individuals who reported that they had a NVQ or HND or HNC degree, individuals who reported other qualifications, and individuals who preferred not to answer were excluded from analyses. The reason for these exclusions was as follows: the first two categories would correspond to 15 and 19 years of education according to the ISCED coding; regarding their mean scores on cognitive ability tests, this might not be the right place for these two degree levels in the ordered hierarchy of educational attainments (Supplementary Figure 1). For the current study, years of education was used a proxy phenotype for cognitive ability ^4,6,8^. A total of 97,550 individuals had valid data for the years of education variable.

**Table 1.**
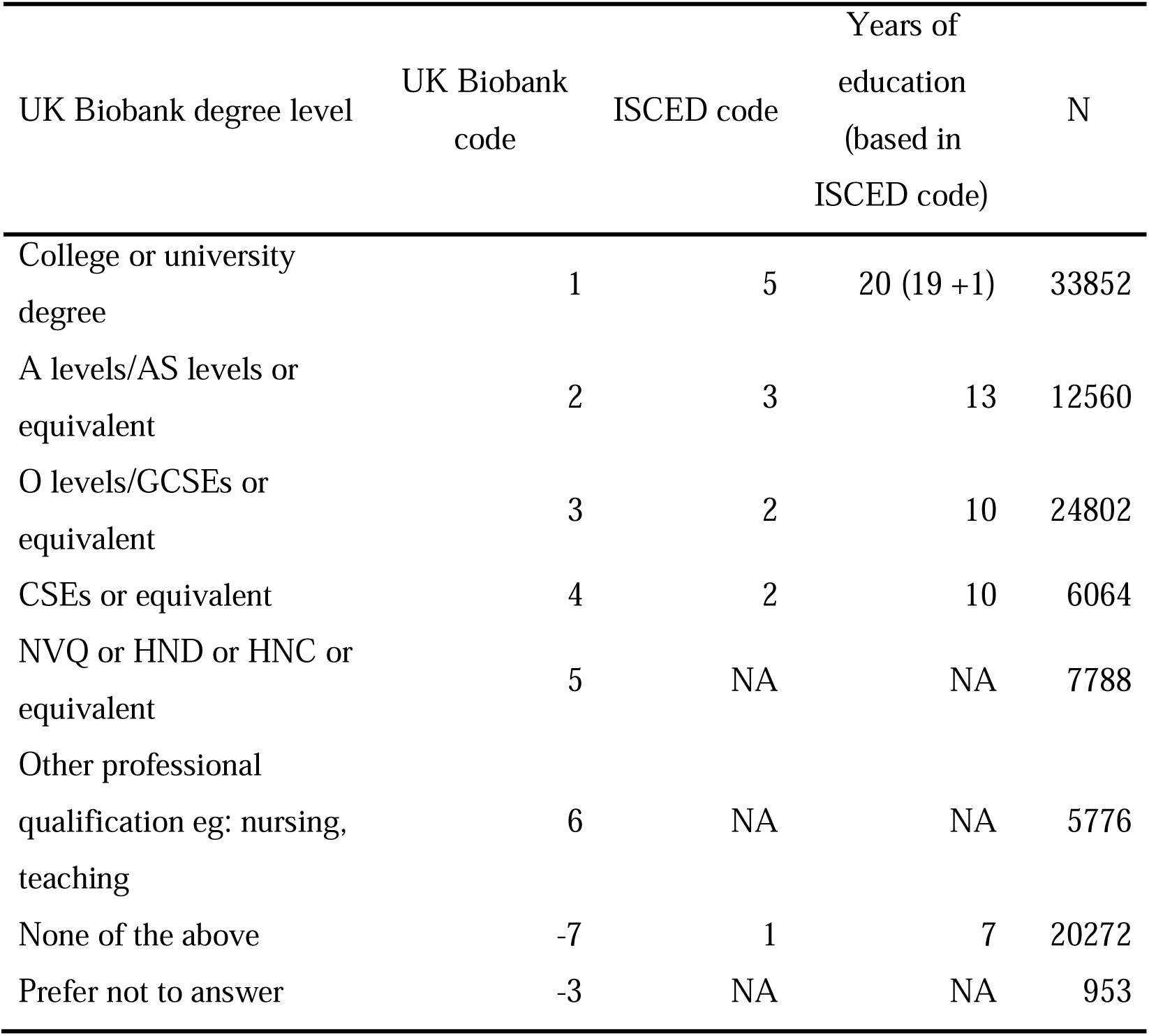
Coding for years of education in UK Biobank based on the ISCED coding^29^ ISCED, 1997 International Standard Classification of Education of the United Nations Educational, Scientific and Cultural Organization.

| UK Biobank degree level | UK Biobank code | ISCED code | Years of education (based in ISCED code) | N |
| --- | --- | --- | --- | --- |
| College or university degree | 1 | 5 | 20 (19 +1) | 33852 |
| A levels/AS levels or equivalent | 2 | 3 | 13 | 12560 |
| O levels/GCSEs or equivalent | 3 | 2 | 10 | 24802 |
| CSEs or equivalent | 4 | 2 | 10 | 6064 |
| NVQ or HND or HNC or equivalent | 5 | NA | NA | 7788 |
| Other professional qualification eg: nursing, teaching | 6 | NA | NA | 5776 |
| None of the above | -7 | 1 | 7 | 20272 |
| Prefer not to answer | -3 | NA | NA | 953 |

### Cognitive ability

Cognitive ability was measured using a 13-item touchscreen computerized verbal-numerical reasoning test. The test included six verbal and seven numerical questions, all with multiple-choice answers, with a two-minute time limit. An example verbal item is: ‘If some flinks are plinks and some plinks are stinks then some flinks are definitely stinks?’ (possible answers: ‘True/False/Neither-true-nor-false/do not know/prefer not to answer’). An example numerical item is: ‘If sixty is more than half of seventy-five, multiply twenty-three by three. If not subtract 15 from eighty-five. Is the answer?’ (possible answers: ‘68/69/70/71/72/do not know/prefer not to answer’). The cognitive ability score was the total score out of 13 (further detail can be found in Hagenaars, et al. ^4^). This test was introduced at a later stage during baseline assessment and only a subset of individuals therefore completed this test. A total of 36 035 had valid cognitive ability and genetic data.

## Covariates

All analyses were adjusted for the following covariates: age when attending assessment centre, sex, genetic batch and array, and the first ten genetic principal components for population stratification.

## Instrumental variables

SNPs associated with each of the five health outcomes and educational attainment were retrieved from the largest available GWAS in European samples for the variables of interest (BMI ^30^, height ^28^, systolic blood pressure ^31^, coronary artery disease ^32^, type 2 diabetes ^33^, and educational attainment ^34^). For educational attainment, we downloaded the summary statistics based on the discovery GWAS only, which did not include the UK Biobank sample. Corresponding SNPs used in the instrumental variables were then extracted from the imputed UK Biobank’s interim release of genotypes, which amounted to 112 151 individuals of self-reported White British ancestry after quality control. Details on the quality control process have been published previously ^4^. SNPs out of Hardy-Weinberg equilibrium (HWE, p < 1×10^−6^), with an imputation quality below 0.9, or individual genotypes with a genotype probability below 0.9 and strand ambiguous SNPs were excluded from the instrumental variables. The individual variants were recoded as 0, 1 or 2 according to the number of trait increasing alleles. Table 2 includes information on the number of SNPs included, and the reference paper. Supplementary Table 3a-f provides details of the included SNPs.

**Table 2.**
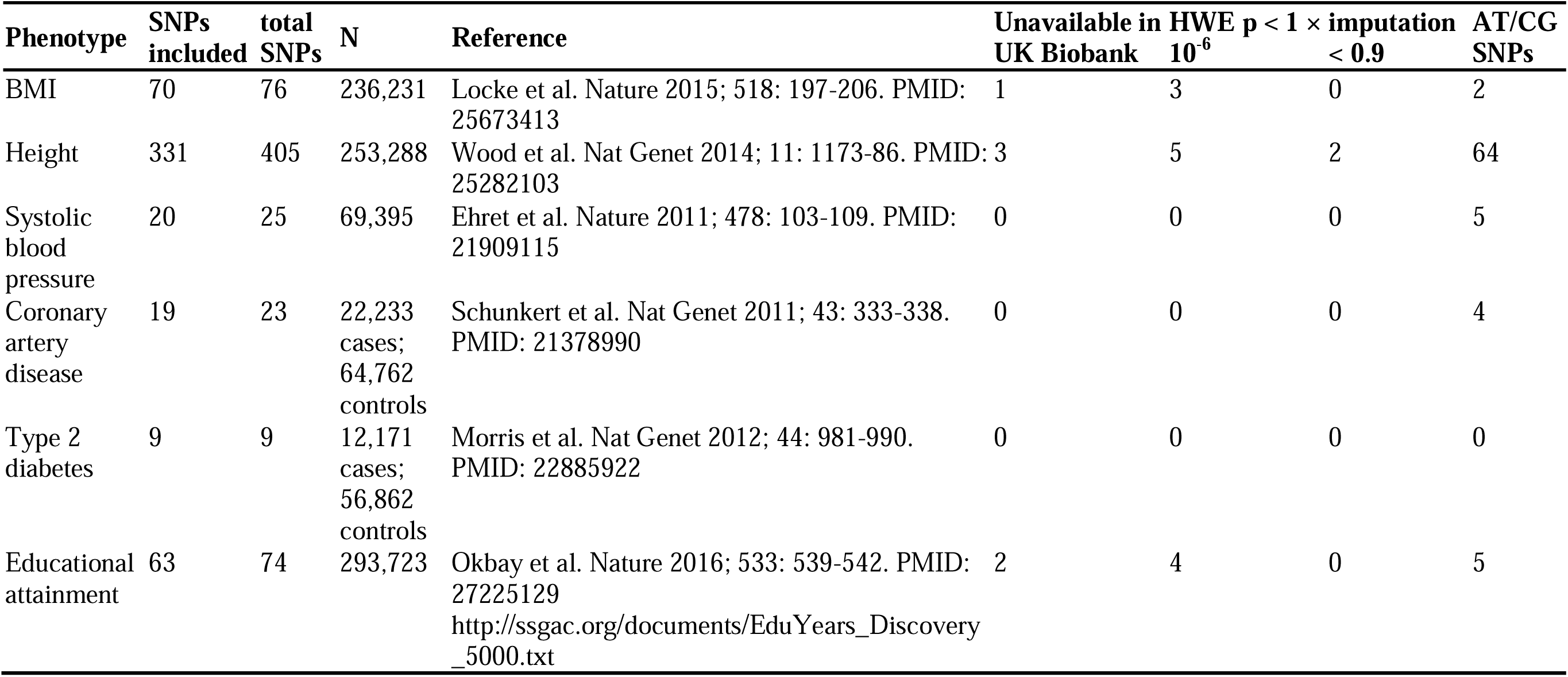
Information about instrumental variables.

**Table 3.** Phenotypic and genetic associations, using Mendelian randomization analysis, between five health instrumental variables and cognitive ability, using the verbal-numerical reasoning test. Significant associations are in bold. OR, odds ratio; MR-IVW, Mendelian randomization - inverse variance weighted method.

| Cognitive ability<br>SNPs (nr) | Phenotypic: health outcomes – cognitive ability |  |  | MR-IVW: health SNPs – cognitive ability |  |  |
| --- | --- | --- | --- | --- | --- | --- |
|  | Beta | 95% CI | p | Beta | 95% CI | p |
| BMI (70) | -0.049 | -0.059 – (-0.039) | **1.51×10<sup>-20</sup>** | -0.035 | -0.147 – 0.077 | 0.5439 |
| Height (331) | 0.1816 | 0.166 - 0.197 | **5.53×10<sup>-124</sup>** | 0.026 | -0.009 – 0.061 | 0.1329 |
| Systolic blood pressure (20) | -0.0492 | -0.061 – (-0.037) | **2.24×10<sup>-17</sup>** | -0.002 | -0.010 – 0.006 | 0.6355 |
| Coronary artery disease (19) | -0.2651 | -0.316 – (-0.214) | **4.62×10<sup>-25</sup>** | -0.018 | -0.045 – 0.009 | 0.2343 |
| Type 2 diabetes (9) | -0.0634 | -0.120 – (-0.007) | **0.0292** | 0.010 | -0.019 – 0.039 | 0.5316 |

## Statistical analysis

### Phenotypic associations

We performed linear regression analysis using BMI, height, systolic blood pressure, coronary artery disease, and type 2 diabetes to predict cognitive ability. We regressed BMI, height, and systolic blood pressure against educational attainment in a linear regression model; coronary artery disease and type 2 diabetes were regressed against educational attainment in logistic regression models.

### Mendelian randomization analysis

The Mendelian randomization analysis was performed using inverse variance weighted regression analysis based on SNP level data, with each instrumental variable (IV) consisting of multiple SNPs ^25^. The inverse variance weighted method is based on a regression of two vectors with the intercept constrained to zero, i.e. the genetic variant with the exposure association, and the genetic variant with the outcome association (Figure 1). By constraining the intercept to zero, this method assumes that all variants are valid instrumental variables based on the Mendelian randomization assumptions. We performed an association analysis between each SNP in the instrumental variable for the exposure and the exposure itself (IV - exposure), as well as between the instrumental variable for the exposure and the outcome (IV - outcome). We then used the vector of the instrumental variable-outcome association analyses against the vector of the instrumental variable-exposure analyses. This association (vector IV - outcome ∼ vector IV - exposure) was weighted by the standard error of the original IV-outcome association, to correct for minor allele frequency, as described by Bowden, et al. ^25^. Power calculations for the MR analyses can be found in Supplementary Table 1. No sensitivity analyses were performed due to the lack of significant causal associations.

## Results

### Health outcomes predicting cognitive ability

BMI, height, systolic blood pressure, and coronary artery disease predicted performance on the verbal-numerical reasoning test of cognitive ability (Table 3). A 1 SD higher BMI was associated with a 0.05 SD lower score for cognitive ability [β = −0.05, 95% CI = −0.06 − (-0.04)]. A 1 SD greater height was associated with a 0.18 SD higher score for cognitive ability [β = 0.18, 95% CI = 0.17 − 0.20]. A 1 SD higher systolic blood pressure was associated with a 0.05 SD lower score for cognitive ability [β = −0.05, 95% CI = −0.06 − (−0.04)]. Individuals with coronary artery disease had, on average, a 0.27 SD lower score for cognitive ability [β = −0.27, 95% CI = −0.32 − (−0.21)]. Individuals with type 2 diabetes had, on average, a 0.06 SD lower score for cognitive ability [β = −0.06, 95% CI = − 0.12 − 0.01]. The Mendelian randomization inverse variance weighted analyses, with the five health outcomes as the exposures, and cognitive ability as the outcome, did not provide any causal evidence for any of these associations.

### Education predicting health outcomes

Educational attainment, as measured by years of education, predicted BMI, height, systolic blood pressure, type 2 diabetes and coronary artery disease (Table 4). The difference between 7 and 20 years of education was associated with a 0.37 SD lower BMI [β = −0.37, 95% CI = −0.39 − (−0.35)], 0.31 SD taller stature [β = 0.31, 95% CI = 0.30 − 0.32], 0.20 lower SBP [β = −0.20, 95% CI = −0.22 − (−0.19)], 0.58 lower odds of type 2 diabetes [OR = 0.58, 95% CI = 0.52 − 0.64], and 0.40 lower odds of coronary artery disease [OR = 0.40, 95% CI 0.37 − 0.43]. The differences between the other groups (7 versus 10 and 13 years of education) can be found in Supplementary Table 2. In every case, the Mendelian randomization inverse variance weighted method did not show a causal effect of educational attainment on the health outcomes. The full results can be found in Table 4.

**Table 4.** Phenotypic and genetic associations, using Mendelian randomization analysis, between the educational attainment instrumental variable and five health outcomes. Significant associations are in bold. OR, odds ratio; MR-IVW, Mendelian randomization - inverse variance weighted method; MR-Egger, Mendelian randomization - Egger regression.

|  | Educational attainment – health outcomes |  |  | Educational attainment SNPs – health outcomes<br>(63 SNPs) |  |  |
| --- | --- | --- | --- | --- | --- | --- |
|  | Phenotypic (7 vs 20 years of education) |  |  | MR-IVW |  |  |
|  | Beta | 95% CI | p | Beta | 95% CI | p |
| BMI | -0.367 | -0.385 – (-0.349) | **<1.00×10<sup>-130</sup>** | -0.026 | -0.157 – 0.105 | 0.6986 |
| Height | 0.312 | 0.300 – 0.324 | **<1.00×10<sup>-130</sup>** | 0.021 | -0.106 – 0.148 | 0.7479 |
| Systolic blood pressure | -0.204 | -0.222 – (-0.187) | **5.61×10<sup>-118</sup>** | -0.003 | -0.097 – 0.091 | 0.9493 |
| Type 2 diabetes | OR: 0.575 | 0.522 – 0.635 | **2.41×10<sup>-28</sup>** | 0.022 | -0.703 – 0.747 | 0.5642 |
| Coronary artery disease | OR: 0.397 | 0.365 – 0.432 | **3.73×10<sup>-101</sup>** | 0.015 | -0.047 – 0.077 | 0.6440 |

## Discussion

This study was designed to investigate causes of the well replicated finding that lower cognitive ability is associated with poorer health outcomes ^1–3^. It used a bidirectional two-sample MR approach to investigate this. We found no evidence for causal association between several health outcomes and cognitive ability, in middle and older age, or between educational attainment and physical health.

Tyrrell, et al. ^13^ showed a significant causal association between taller stature and time spent in full time education in UK Biobank. They did not find a causal association between taller stature and degree level. The measure of time spent in full time education in UK Biobank excluded individuals who reported having a college degree, which could explain the discrepancy in results. The current study did include individuals who reported having a college degree, however used a categorical measure of four categories, whereas Tyrrell, et al. ^13^ used a continuous measure of time spent in full time education. In a non-peer-reviewed (at the time of writing) study, Tillmann, et al. ^35^ did report a causal association from educational attainment to coronary artery disease and BMI using a two-sample MR approach based on two independent consortia^35^. They used data from two independent GWAS consortia, including 349,306 individuals for educational attainment, 194,427 (63,746 cases) individuals for coronary artery disease, and 339,224 individuals for BMI. The current study used the same data for educational attainment on a subset of individuals (N = 293,723), and 111,712 individuals with BMI data; however, coronary artery disease was based on self-report diagnosis in UK Biobank, which included 110,072 (5288 cases) individuals. The summary level data for coronary artery disease in the Tillmann, et al. ^35^ report included both European and East-Asian individuals, whereas the current study only includes individuals of White British ancestry. They^35^ excluded overlapping cohorts between educational attainment and coronary artery disease data; however, it is unclear if overlapping cohorts were excluded for BMI.

Another explanation for the lack of causal associations in the present study could be the high polygenic aetiology of the traits analysed in this study. Instrumental variables for cardiovascular disease, type 2 diabetes, blood pressure, and educational attainment explain a small amount of the variance in the exposure. A better instrumental variable would be expected to explain a substantial amount of the variance of the exposure. As shown by the power calculations (Supplementary Table 1), all instrumental variables (except BMI and systolic blood pressure) had sufficient power to detect the same magnitude of association as the observational estimates. The low power for BMI and systolic blood pressure potentially explains the lack of association with cognitive ability. A previous study by the current authors indicated a degree of genetic overlap between cognitive ability and health across the genome ^4^ The idea of genetic overlap between health and cognitive ability is consistent with the theoretical construct of bodily system integrity ^24^, whereby a latent trait is manifest as individual differences in how effectively people meet cognitive and health challenges from the environment, and which has some genetic aetiology.

Strengths of this study include the large sample size of UK Biobank, the participants of which all took the same cognitive tests, completed the same questionnaires and answered the same interview questions, in contrast to most genetic studies, where assessments across different cohorts often vary. A further strength is the fact that all of the UK Biobank genetic data were processed in a consistent matter, on the same platform and at the same location. The genetic variants on which the instrumental variables originated used the largest available GWAS at moment of testing.

Limitations of this study include the fact that cognitive ability was only measured on a subset of the UK Biobank participants and that it was a bespoke test. A second major limitation was that there is no published large genome-wide association study of cognitive ability in early life from which we could obtain genetic variants to use as an instrumental variable. Therefore, we used genome-wide significant SNPs associated with educational attainment as our early life cognitive ability instrument. A further limitation is the case-control ascertainment in UK Biobank, as the current study based case-control status on self-report measures. This may have led to misclassification of disease status, causing a likely bias towards the null hypothesis^36^.

Overall, this study found phenotypic cognitive-physical health associations, but did not find evidence for causal associations between cognitive ability and physical health. This may be due to weak instrumental variables, poorly measured outcomes, or the small numbers of disease cases. Future work should therefore focus on stronger instrumental variables, as well as better measurement of the outcome variables.

## Acknowledgements

This work was supported by The University of Edinburgh Centre for Cognitive Ageing and Cognitive Epidemiology, part of the cross council Lifelong Health and Wellbeing Initiative (MR/K026992/1). Funding from the Biotechnology and Biological Sciences Research Council (BBSRC) and Medical Research Council (MRC) is gratefully acknowledged. This research was conducted using the UK Biobank Resource.

## Author contributions

All authors contributed to the design of the study. UK Biobank collected all data and was involved in pre-processing the genetic data. SPH extracted, pre-processed, and analysed the data. SPH and SEH wrote the main manuscript text. All authors discussed and commented on the manuscript.

